# The contribution of cross-modal sensory input to reaching and grasping in blind and sighted short-tailed opossums (*Monodelphis domestica*)

**DOI:** 10.64898/2026.05.25.727491

**Authors:** Carlos R Pineda, Faith Zirkelbach-Ngai, Madeleine Miller, Leah A Krubitzer

## Abstract

Reaching and grasping are essential goal-directed behaviors that require the integration of visual, olfactory, and somatosensory inputs. Although the loss of vision can profoundly disrupt sensory-guided behaviors, mammals often exhibit compensatory cross-modal plasticity allowing them to reach and grasp with ease. To examine how early sensory loss shapes reaching behavior, we performed bilateral enucleations in short-tailed opossums (*Monodelphis domestica*) of either sex at postnatal day 4, before retino-thalamic and thalamocortical connections have formed. We assessed performance in early blind (EB) and sighted control (SC) animals using a semi-naturalistic reach-to-grasp task requiring precise unilateral limb targeting to retrieve a dead cricket. To isolate the contributions of specific sensory modalities to this task, we selectively disrupted olfactory and mystacial vibrissae inputs and manipulated lighting conditions during task performance. EB opossums were capable of accurate reaching and grasping and showed no significant differences in performance relative to SC animals. Both groups relied strongly on tactile input, as whisker trimming significantly increased targeting error. Removal of olfactory input also impaired performance, with a disproportionately greater effect in EB animals. These findings show that short-tailed opossums retain functional reach-to-grasp behavior after early vision loss and that accurate forelimb movements arise from enhanced spared sensory systems.

**Significance Statement:** Congenital sensory loss in humans alters the sensory landscape used to navigate the world, making compensatory strategies mediated by the spared sensory systems essential for goal-directed behaviors such as reaching and grasping. Although these behaviors have been widely studied across species, the extent to which spared senses support their execution after congenital vision loss remains unclear. Here, we use the short-tailed opossum as a model of congenital blindness to quantify the contributions of whisker-mediated touch and olfaction to reaching performance. We show that both early blind and sighted opossums rely on whisker touch for reaching and grasping, but that olfaction plays a profound role in task performance in early blind opossums.

## Introduction

Reaching and grasping is an ethologically relevant behavior and demands multimodal sensory processing and sensorimotor integration. While most studies that examine these behaviors are performed in non-human primates, reaching and grasping is a ubiquitous behavior for most mammals with well-developed forelimbs and forepaws. Although scientists often focus on understanding how a single sensory system contributes to species-specific behaviors, in natural environments animals rely on the integration of multiple sensory modalities in a context-dependent manner to generate adaptive behavior. Effective motor control likely depends on the integration of visual, auditory, proprioceptive, tactile, vestibular and even olfactory inputs that contribute to different stages of movement planning and execution (Stone and Gonzalez, 2015).

Because of the importance of multimodal processing, loss of one sensory modality, whether congenital or acquired, profoundly alters sensory experience and can disrupt normal behaviors. Even when faced with such disruptions, mammals exhibit remarkable flexibility, compensating both neurophysiologically and behaviorally, particularly when sensory loss occurs during early development (Kahn and Krubitzer, 2002; Karlen et al., 2006; Fujii et al., 2009; Wong et al., 2011a; Englund et al., 2020; Ramamurthy et al., 2021; Asumbisa et al., 2022).

Studies in humans, non-human primates, ferrets, mice, rats, and hamsters have demonstrated that the effects of early sensory loss on cortical organization vary substantially across cortical areas and species. Studies in non-primate mammals often describe extensive reorganization of primary sensory cortices, whereas studies in primates report more variable effects across cortical regions (Sharma et al., 2000; Izraeli et al., 2002; Chabot et al., 2008; Voller et al., 2014; Hagan et al., 2017; Wang et al., 2017; Setti et al., 2023; Maruoka et al., 2024). However, most of these studies examined sensory loss at relatively late stages of development when anatomical pathways are partially or completely developed.

**Table 1:** List of Abbreviations.

| Abbreviation | Definition |
| --- | --- |
| EB | Early Blind |
| EMM | Estimated Marginal Means |
| hMT+ | Human Middle Temporal Area |
| P4 | Postnatal Day 4 |
| S1 | Primary Somatosensory Area |
| SC | Sighted Control |
| SPC | Superior Parietal Cortex |
| V1 | Primary Visual Area |
| V2 | Second Visual Area |
| V3 | Third Visual Area |

In the current study, we examine the effects of early loss of vision on reaching behavior in the short-tailed opossum (*Monodelphis domestica*), and the contribution of tactile and olfactory inputs on task performance. The short-tailed opossum is precocial, allowing us to remove visual inputs via bilateral enucleations before retino-thalamic and thalamo-cortical connections have formed (postnatal day 4; P4), thereby approximating a model of congenital anophthalmia. Previous studies in our laboratory demonstrate that these early enucleations lead to changes in the connectivity and response properties of neurons in the putative visual and somatosensory cortex (Kahn and Krubitzer, 2002; Karlen et al., 2006; Ramamurthy and Krubitzer, 2018). For example, neurons in the primary visual cortex (V1) in early blind (EB) short-tailed opossums respond to auditory and tactile stimulation (Figure 1A), and neurons in the whisker representation in primary somatosensory cortex (S1) undergo receptive field modifications that may subserve the greater texture discrimination abilities observed in blind animals (Kahn and Krubitzer, 2002; Ramamurthy and Krubitzer, 2018; Ramamurthy et al., 2021; Figure 1B,C). Interestingly, EB opossums also outperform SC counterparts on tasks where tactile input from the whiskers compensates for the lack of vision (Englund et al., 2020; Ramamurthy et al., 2021).

**Figure 1:**
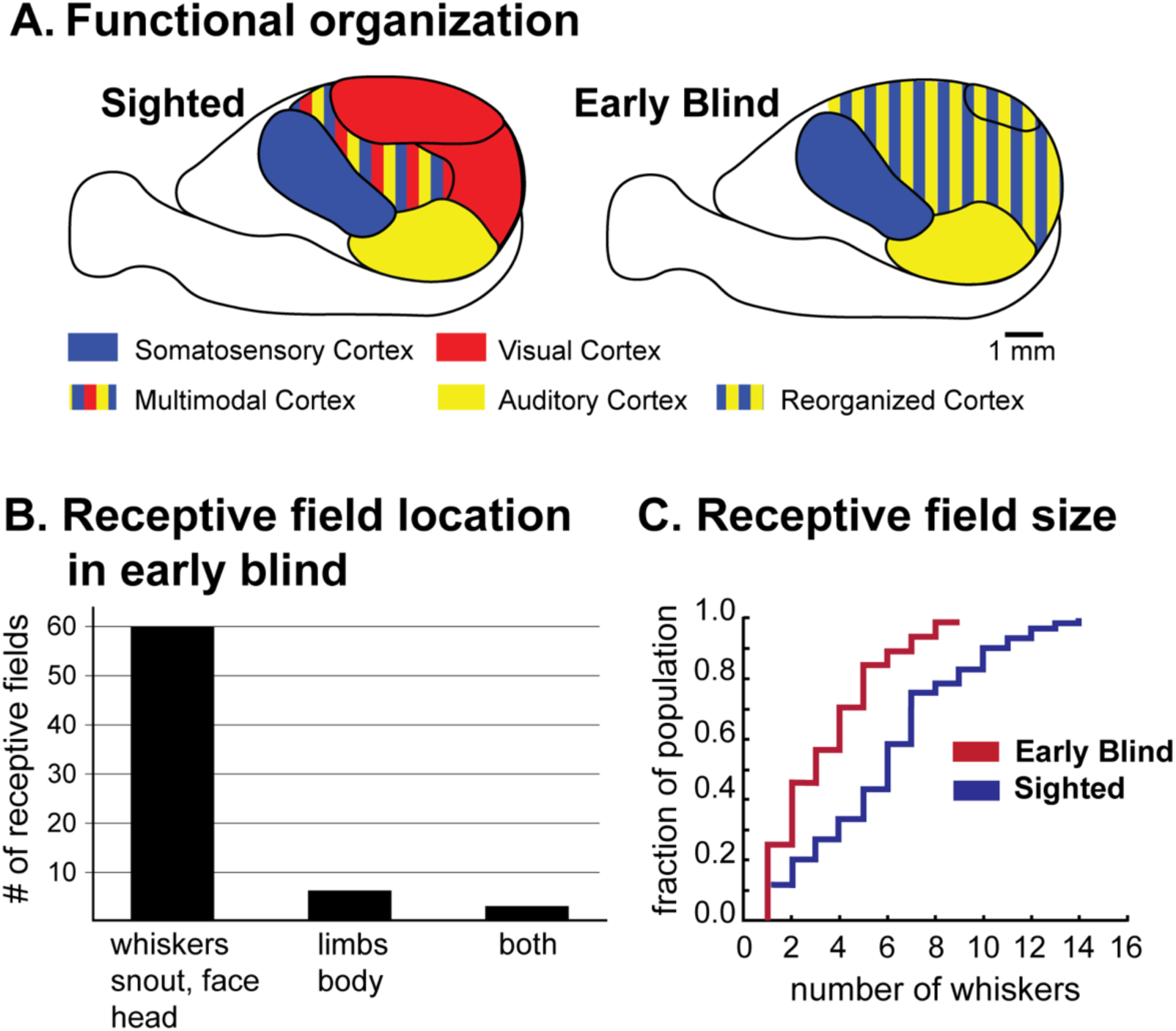
Alterations in the functional organization of V1 and receptive fields for neurons in V1 and S1 following early loss of vision. (**A**) The organization of neocortex in sighted (left) and early blind opossums (right). The neurons in re-organized V1 and other adjacent visual areas of early blind opossums respond to auditory and tactile stimulation. (**B**) Bar graph showing the proportion of neurons in re-organized V1 and other “visual areas” with receptive fields representing the face, head, and lower extremities of early blind short-tailed opossums. (**C**) Line graph showing the reduction in receptive field size of neurons in the whisker representation of S1 in early blind (red) and sighted control (blue) short-tailed opossums. Adapted from Khan et al, 2002 (A and B) and Ramamurthy et al, 2019 (**C**).

We assessed whether EB short-tailed opossums compensate for lack of vision using their remaining sensory modalities by examining their performance on a semi-naturalistic and widely used reach-to-grasp task that requires precise sensory-guided targeting of the forelimb and paw to retrieve a reward (Ivanco et al., 1996; Whishaw, 1996; Klein et al., 2012; Whishaw et al., 2017). To determine the contribution of individual sensory modalities to reach targeting after vision loss, we performed a series of experiments in which we selectively and reversibly removed input from individual sensory modalities. These manipulations included extinguishing light sources (for SC), trimming mystacial vibrissae, and temporarily inducing anosmia. To our knowledge, this is the first investigation to examine the contribution of multiple sensory inputs to reaching and grasping behavior.

## Methods

Reaching behavior was examined in 21 adult (>180 days old) short-tailed opossums (*Monodelphis domestica*) weighing 70 to 120 grams. Eleven animals (7 females, 4 males) were bilaterally enucleated (EB) at postnatal day 4 (P4), and 10 animals (7 females, 3 males) served as sighted littermate controls (SC). All animals were born and housed in the UC Davis Psychology Department vivarium under a 14:10 hour light/dark cycle (lights on at 6 AM), with food and water available ad libitum. Animals were weaned at 56 days and separated from littermates at four months. All procedures were approved by the UC Davis Institutional Animal Care and Use Committee and followed NIH guidelines.

### Bilateral Enucleation Surgery

Bilateral enucleations were performed at P4 as described previously (Ramamurthy et al, 2018). Briefly, dams were anesthetized with isoflurane (5%) and maintained under Alfaxalone (3.0 mg/kg IM). Pups were anesthetized using hypothermia. The skin overlying the developing eye was incised, the immature retina and optic tissue removed, and the socket flushed with sterile saline (9%). The incision was sealed with surgical glue (GLUture, Zoetis Inc. Kalamazoo MI). Half the litter was enucleated, while the remaining pups served as sighted controls. Dams recovered in their home cage following IACUC guidelines.

### Behavioral Testing

Adult EB and SC opossums were trained for five days to perform a reach-to-grasp task. The reaching box was modeled based on that described previously for rodents (Whishaw et al., 2010; Alaverdashvili and Whishaw, 2013; Figure 2A). The box was made of clear Plexiglass (350 x 250 x 300 mm). A vertical slit centered in the front of the box (10 mm wide, 150 mm tall) served as an opening through which opossums could reach for a dead cricket reward placed on a shelf attached to the outside of the reaching box. This shelf was positioned 40 mm above the floor of the box, and the cricket was placed on a well in the shelf 20 mm away from the slit opening (Figure 2B).

**Figure 2:**
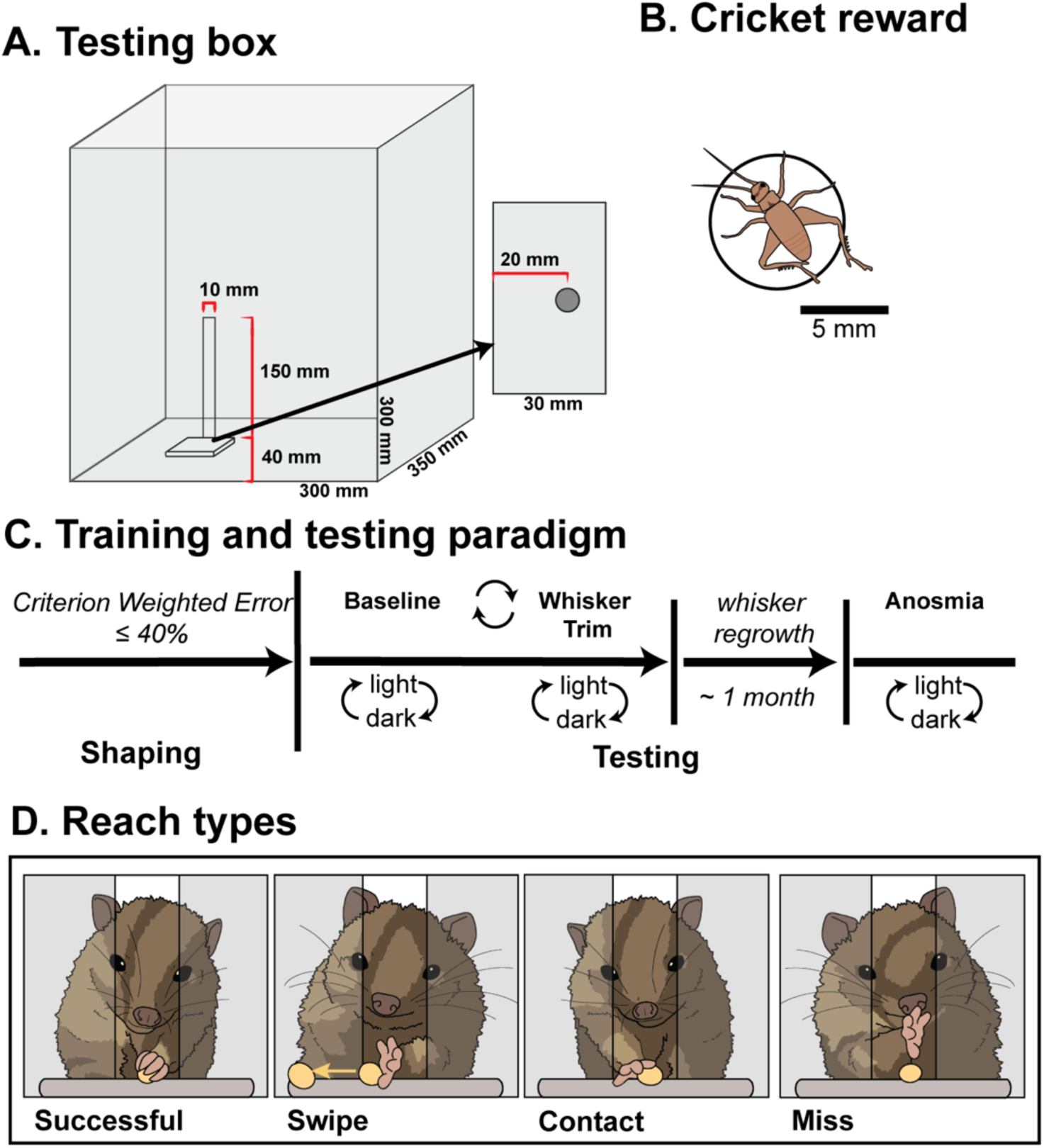
Reaching and grasping test box and testing paradigm for the short-tailed opossum. **(A)** Front view of the reaching and grasping box constructed for short-tailed opossums. Animals were trained to reach through a slit for a cricket reward placed on a platform approximately 20 mm away from the slit opening. The cricket reward was positioned in a central well to ensure it remained in place. The dimensions of the box are shown in the schematic. **(B)** Image depicting the size and shape of the cricket reward relative to well size (circle around cricket. **(C)** Animal training and testing paradigm. Animals were shaped for five days and were tested after achieving a weighted error below 40%. Once criterion performance was achieved, animals were tested under light and dark conditions with whiskers intact (baseline), whisker-trimmed, and anosmic. To minimize potential order effects associated with task learning, the sequence of baseline and whisker-trimmed testing, as well as light and dark testing conditions, was counterbalanced across animals. **(D)** Schematic illustrations depicting the four reach outcome categories scored during the task: Successful, Swipe, Contact, and Miss.

Training consisted of placing short-tailed opossums inside the reaching box for 10-minute sessions, during which dead crickets were placed around the slit and on the platform. All animals underwent a standardized shaping procedure to learn the association between the slit, platform and reward, although some animals more readily initiated reaching behavior than others. During shaping, crickets were initially positioned near or within the slit and were gradually moved to the target indentation on the platform as animals learned to associate the slit with reward delivery. As learning progressed, cricket placements were restricted to the platform, and the distance from the slit was increased to shape reaching behavior. To discourage repeated reaching attempts for a single reward, which were common, animals were limited to one attempt per trial by alternating placement of the target cricket on the platform with a second reward placed at the rear of the testing chamber. Animals were considered shaped once they reliably reached for crickets placed on the target indentation, regardless of success rate. Following shaping, animals underwent 5 days of training, and performance was evaluated. Animals that did not reach performance criterion below 40% weighted error within 5 days were excluded from further testing.

Reach-to-grasp tests consisted of 20-30 trials per session in which 5-10 mm-sized dead crickets (Figure 2B) were presented over a maximum period of 10 minutes. Because of individual variability in reach initiation (See Table 2), each animal completed two testing sessions under each experimental condition. Data collection began after 5 days of training. Each <u>Visual Group (EB, SC)</u> was tested under three <u>Sensory Conditions</u>: baseline, whisker-trimmed, and anosmic. The order of baseline and whisker-trimmed testing was counterbalanced across animals. Testing was also conducted under both light and dark conditions (Lighting Condition), which were counterbalanced across consecutive days (Figure 2C). A 660 nm wavelength red lamp was used to provide visibility for experimenters during trials conducted in darkness. Previous studies from our laboratory show that performance under this illumination does not differ from complete darkness (Englund et al., 2020), and it is known that short-tailed opossums possess long-wavelength sensitive (LWS) cones with an expected peak sensitivity in the ∼500 – 570 range (Hunt et al., 2009), placing 600 nm well outside this range. However, this does not completely rule out residual photoreceptor activation under these conditions.

**Table 2:** Number of testing sessions contributed by each animal under each experimental condition. Values indicate the number of testing sessions included in the behavioral analyses.

| Animal | Sex | Baseline<br>Light | Baseline<br>Dark | Trimmed<br>Light | Trimmed<br>Dark | Sham | Hab./Dishab. | Anosmia<br>Light | Anosmia<br>Dark |
| --- | --- | --- | --- | --- | --- | --- | --- | --- | --- |
| SC1 | F | 2 | 1 | NA<sup>†</sup>A | NA<sup>†</sup>A | NA<sup>†</sup>A | NA<sup>†</sup>A | NA<sup>†</sup>A | NA<sup>†</sup>A |
| SC2 | F | 1 | 2 | 2 | 0<sup>†</sup>C | 1 | 1 | 2 | 1 |
| SC3 | F | 2 | 2 | 2 | 2 | 1 | 1 | 0<sup>†</sup>C | 1 |
| SC4 | F | 1 | 0<sup>†</sup>C | 2 | 1 | 1 | 1 | 1 | 1 |
| SC5 | F | 1 | 1 | 2 | 1 | 1 | 1 | 1 | 1 |
| SC6 | F | 2 | 1 | 0<sup>†</sup>C | 0<sup>†</sup>C | 1 | 1 | 0<sup>†</sup>C | 0<sup>†</sup>C |
| SC7 | F | 1 | 1 | NA<sup>†</sup>A | NA<sup>†</sup>A | NA<sup>†</sup>A | NA<sup>†</sup>A | NA<sup>†</sup>A | NA<sup>†</sup>A |
| SC8 | M | 1 | 1 | 1 | 1 | 1 | 1 | 1 | 0<sup>†</sup>C |
| SC9 | M | 2 | 1 | 1 | 2 | 1 | 1 | 2 | 2 |
| SC10 | M | 1 | NA<sup>†</sup>A | NA<sup>†</sup>A | NA<sup>†</sup>A | NA<sup>†</sup>A | NA<sup>†</sup>A | NA<sup>†</sup>A | NA<sup>†</sup>A |
| EB1 | F | 1 | 1 | 1 | 1 | NA<sup>†</sup> | 1 | 0<sup>†</sup>C | 0<sup>†</sup>C |
| EB2 | F | 2 | 2 | 1 | 0<sup>†</sup>C | 1 | 1 | 2 | 1 |
| EB3 | F | 1 | 0<sup>†</sup>C | 2 | 1 | 1 | 1 | 1 | 1 |
| EB4 | F | 2 | 2 | 1 | 0<sup>†</sup>C | NA<sup>†</sup> | 1 | 0<sup>†</sup>C | 0<sup>†</sup>C |
| EB5 | F | 2 | 2 | 2 | 0<sup>†</sup>C | NA<sup>†</sup> | 1 | 1 | 1 |
| EB6 | F | 1 | 1 | 1 | 1 | NA<sup>†</sup> | 1 | NA<sup>†</sup>B | NA<sup>†</sup>B |
| EB7 | F | 1 | 1 | 1 | 1 | NA<sup>†</sup> | 1 | NA<sup>†</sup>B | NA<sup>†</sup>B |
| EB8 | M | 2 | 0<sup>†</sup>C | 2 | 1 | NA<sup>†</sup> | 1 | 1 | 0<sup>†</sup>C |
| EB9 | M | 1 | 0<sup>†</sup>C | 1 | 0<sup>†</sup>C | 1 | 1 | 2 | 1 |
| EB10 | M | 1 | 2 | 1 | 2 | 1 | 1 | 0<sup>†</sup>C | 1 |
| EB11 | M | 2 | 1 | 2 | 1 | 1 | 1 | NA<sup>†</sup>B | NA<sup>†</sup>B |

*Abbreviations:* F, female; M, male.
NA<sup>†</sup>, animal not included in this condition.
NA<sup>†</sup>A, animal not tested because of attrition before testing.
NA<sup>†</sup>B, anosmia data excluded because the animal failed anosmia validation.
0<sup>†</sup>C, testing was performed but no sessions contributed because the animal did not perform any reaches.

### Whisker trimming and induction of anosmia in adults

To determine the extent to which other sensory systems contributed to task performance, we first removed facial whiskers and then induced anosmia after allowing whiskers to regrow. For whisker trimming, animals were briefly anesthetized with isoflurane (2–5%), and all mystacial, submandibular, and genal vibrissae were trimmed to ∼1 mm in length. Following whisker trimming, behavioral testing was done for two consecutive days. Whiskers were allowed to grow back (∼30 days), and then anosmia was induced by intranasal infusions of zinc sulfate (50 µL, 5%) into each nostril, administered 1 hour apart. We generated sham controls in a subset of animals (5 blind, 7 control; see Table 2), in which sterile saline (50 µL, 9%) was infused into each nostril under the same anesthetic regimen and at the same time intervals stated above. Animals recovered in their home cage for 10–12 hours before undergoing sham or anosmia reach testing.

In animals that had undergone sham and zinc sulfate infusions, we conducted an olfactory habituation/dishabituation experiment to ensure that anosmia was successfully induced. The task consisted of four vehicle presentation epochs (Vehicle 1 – 4; mineral oil; Thermo Fisher Scientific, Waltham, Massachusetts) followed by four odor exposure epochs (Odor 1 – 4). Each epoch lasted 4 minutes and was separated by a 1-minute intertrial period. The odor of choice was varied across anosmic and sham testing to maximize the novelty effect of each odor. Odors used were hexyl acetate, hexanoic acid, amyl acetate, and ethyl acetate (all 1%, Thermo Fisher Scientific, Waltham, Massachusetts), each diluted in mineral oil.

Three independent observers blind to the animals’ anosmia condition scored habituation/dishabituation task videos. Behavioral scoring was based on a predefined ethogram comprising two categories: (1) direct contact with the odor delivery vehicle and (2) attentive behavior, defined by forward movement, proximity to the vehicle, or odor. Time in direct contact and time attending to the vehicle or odor were summed within each epoch and expressed as a percent of total epoch duration (Equation 1: Time Spent Sniffing (%)). The percentage of time spent sniffing was then averaged across anosmic and sham conditions. Only animals confirmed to be behaviorally anosmic were included in the behavioral analysis (See Table 2). All testing was conducted at least 10 hours post-anesthetic induction and within 72 hours of whisker trimming and anosmia induction procedures.

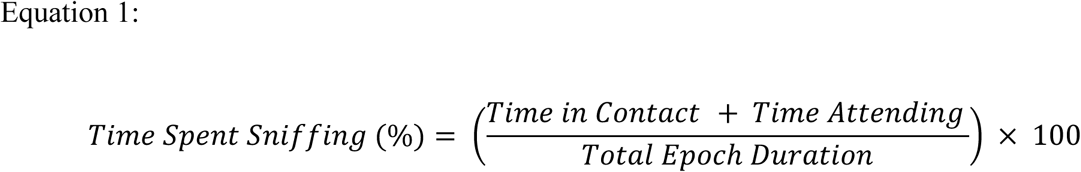

### Video Analysis and Scoring Reaching Behavior

Reaching behavior was recorded using three orthogonally positioned GoPro cameras (240 fps; GoPro, Inc, San Mateo, California). Videos were analyzed with BORIS behavioral scoring software (Friard and Gamba, 2016). Three trained independent observers, blind to the animals’ Sensory Condition, scored all trials. We adapted a four-point scoring system originally developed by Metz and Wishaw (2002). This scoring system includes four categories: <u>Total Miss</u>: 0 points are awarded when the reaching forepaw completely misses the cricket reward; <u>Swipe</u>: 1 point is awarded when the reaching forepaw contacts the cricket reward such that the cricket is knocked off the platform; <u>Contact</u>: 2 points are rewarded when the reaching forepaw contacts the cricket reward but is not successfully grasped and retrieved; <u>Success</u>: 3 points are awarded when the reaching forepaw grasps and retrieves the cricket (Equation 2: Weighted Error Formula; Figure 2D). Although efforts were made to restrict animals to one reach per bout, sometimes many attempts occurred. We only scored the first attempt during each bout.

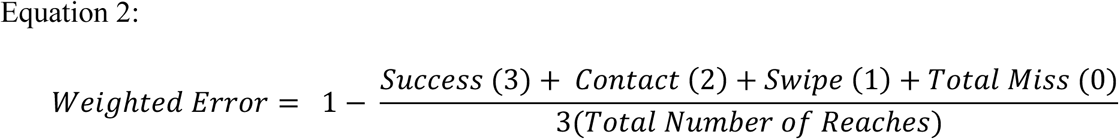

## Statistical Analysis

### Inter-rater reliability

To assess inter-rater reliability, we computed intraclass correlation coefficients (ICC3, single fixed raters) across independent scorers who were blind to the experimental conditions. Reliability was high (ICC3 = ICC3 = 0.96, p < 0.001). Linear regression analysis further demonstrated strong correspondence between scorers (R²adj = 0.868, F(1, 32) = 218.0, p < 0.001; Figure S1).

### Analysis of Habituation/Dishabituation Task

To verify the effectiveness of the anosmia induction protocol, we analyzed sniffing behavior during a habituation/dishabituation task using a repeated-measures ANOVA (statsmodels package in Python (Seabold and Perktold, 2010). Time spent sniffing (%) was compared between pre- and post-anosmia conditions (i.e., pre- and post-induction) within each subject. We used a repeated-measures ANOVA to test for differences within and across testing days and applied the Holm–Sidak method to correct for multiple comparisons. Animals ceased to respond to odor stimuli following anosmia induction (adjusted R² = 0.44, F (5) = 6.33, p = 0.012; pre-anosmia vs post-anosmia Odor 1: Tukey’s HSD: p < 0.0001; Figure S2A). We observed a general decrease in interest in the vehicle when no odor was present after anosmia induction compared to when intact (Vehicle 1: Tukey’s HSD: p = 0.0267). Animals who underwent sham procedures continued to investigate the initial odor presentation at levels significantly greater than post-anosmia animals (Odor 1, Sham vs Post-Anosmia: Tukey’s HSD: *p = 0.03*; Figure S2A), indicating that odor-directed investigatory behavior persisted following sham procedures. Reach-to-grasp performance also differed significantly between sham and anosmic conditions in both SC and EB animals (Paired t-test: t-statistic = -2.707, p = 0.024; S2B), indicating that olfactory deprivation impaired task performance beyond the effects of the infusion procedure alone.

### Analysis of Reaching Data

We initially fit a linear mixed-effects model including all main effects and interactions among Visual Group (EB/SC), Sensory Condition (baseline/whisker trimmed/anosmic), and Lighting Condition (light/dark), along with Biological Sex and a random intercept for Animal ID (See Initial Model). Because each animal was tested repeatedly across sensory and lighting conditions, animal identity (ID) was included as a random intercept to account for within-subject correlations in weighted error (Initial Model). Model simplification was performed using backward hierarchical selection, and higher-order interactions were sequentially removed based on likelihood ratio and Akaike Information Criterion values. The final model retained interactions between Visual Group and Sensory Condition and between Visual Group and Lighting Condition, along with the main effects of these variables and Biological Sex (See Final Model). To specifically evaluate the contribution of visual cues under intact sensory conditions, a separate linear mixed-effects model was fit to baseline trials only. This model included Visual Group, Lighting Condition, their interaction, Biological Sex, and a random intercept for Animal ID, allowing the effect of lighting on baseline reach-to-grasp performance to be assessed independently of the whisker-trimming and anosmia manipulations (See Baseline Model).

Post-hoc pairwise comparisons of estimated marginal means (EMM) were conducted using the emmeans package (Lenth, 2025; version 1.11.1) in R (version 12.1), with Tukey’s correction applied to adjust for multiple comparisons. All contrasts reported are adjusted for biological sex to account for sexual dimorphism. The results of EMM contrasts are found in Table 3.

**Table 3:**
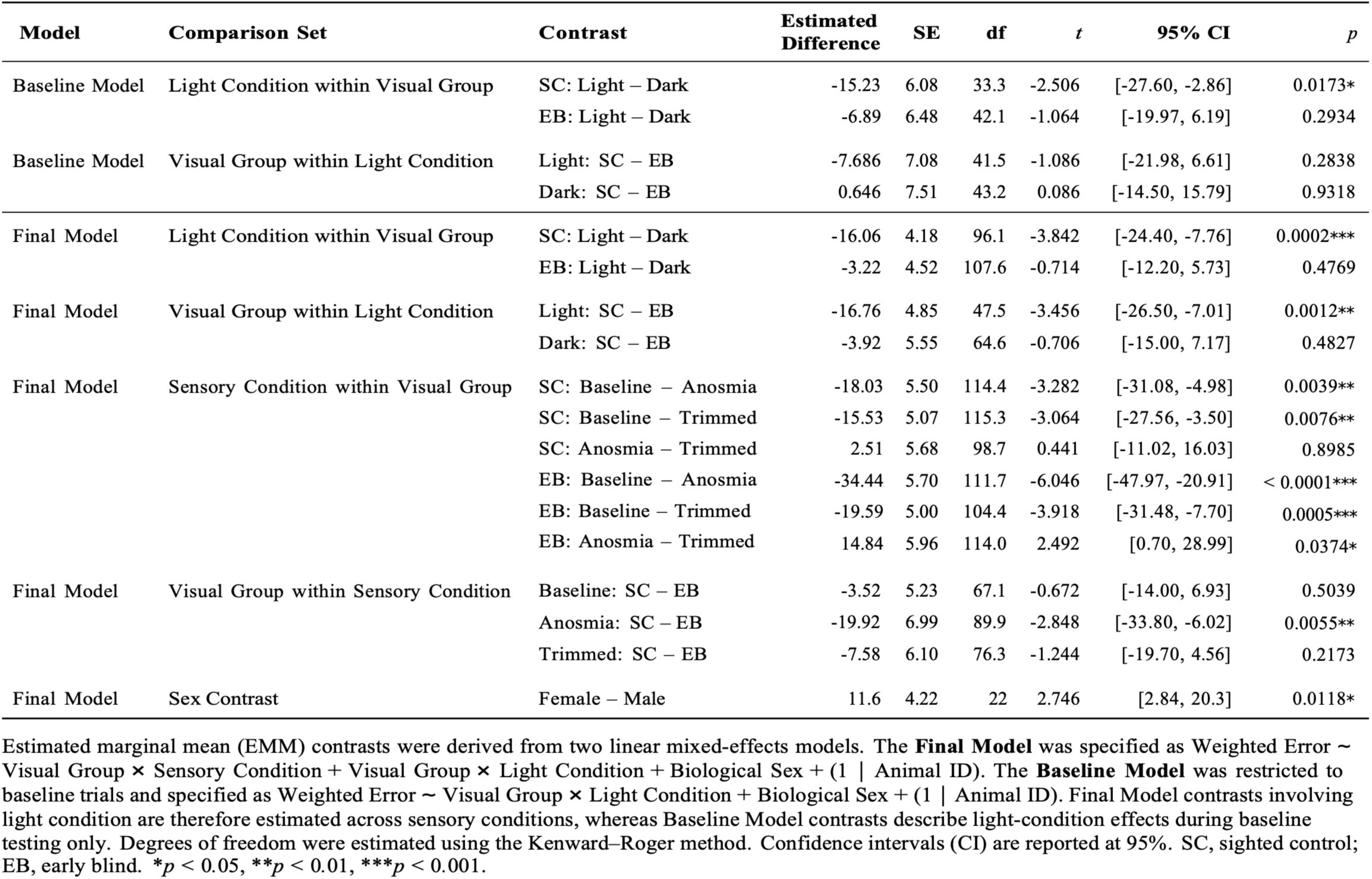
Post hoc contrasts derived from the baseline and final linear mixed-effects models.

All remaining statistical analyses were conducted in Python using custom analysis scripts, including paired t-tests comparing reach-to-grasp errors between sham and anosmic conditions, repeated-measures ANOVA analyzing habituation/dishabituation behavior, Mann-Whitney U tests assessing differences in paw preference across visual conditions, and inter-rater reliability analysis evaluating scoring consistency between observers.

**Initial Model: *Weighted Error ∼ Visual Group × Sensory Condition × Lighting Condition + Biological Sex + (1 | Animal ID)***

**Final Model: *Weighted Error ∼ Visual Group × Sensory Condition + Visual Group × Light Condition + Biological Sex + (1 | Animal ID)***

**Baseline Model: *Weighted Error ∼ Visual Group × Light Condition + Biological Sex + (1 | Animal ID)***

### Code and Data Accessibility

Custom Python scripts used for behavioral analysis are available in GitHub at https://github.com/guatomallin/RGANALYSIS. Data supporting the findings of this study are available from the corresponding author upon reasonable request.

## Results

We trained early blind (EB) and sighted control (SC) adult opossums to perform a reach-to-grasp task (Figure 3). To determine the extent to which EB and SC opossums relied on non-visual cues, we sequentially restricted whisker and olfaction-mediated sensory sampling by whisker trimming, and subsequently by inducing anosmia after the whiskers had regrown.

**Figure 3:**
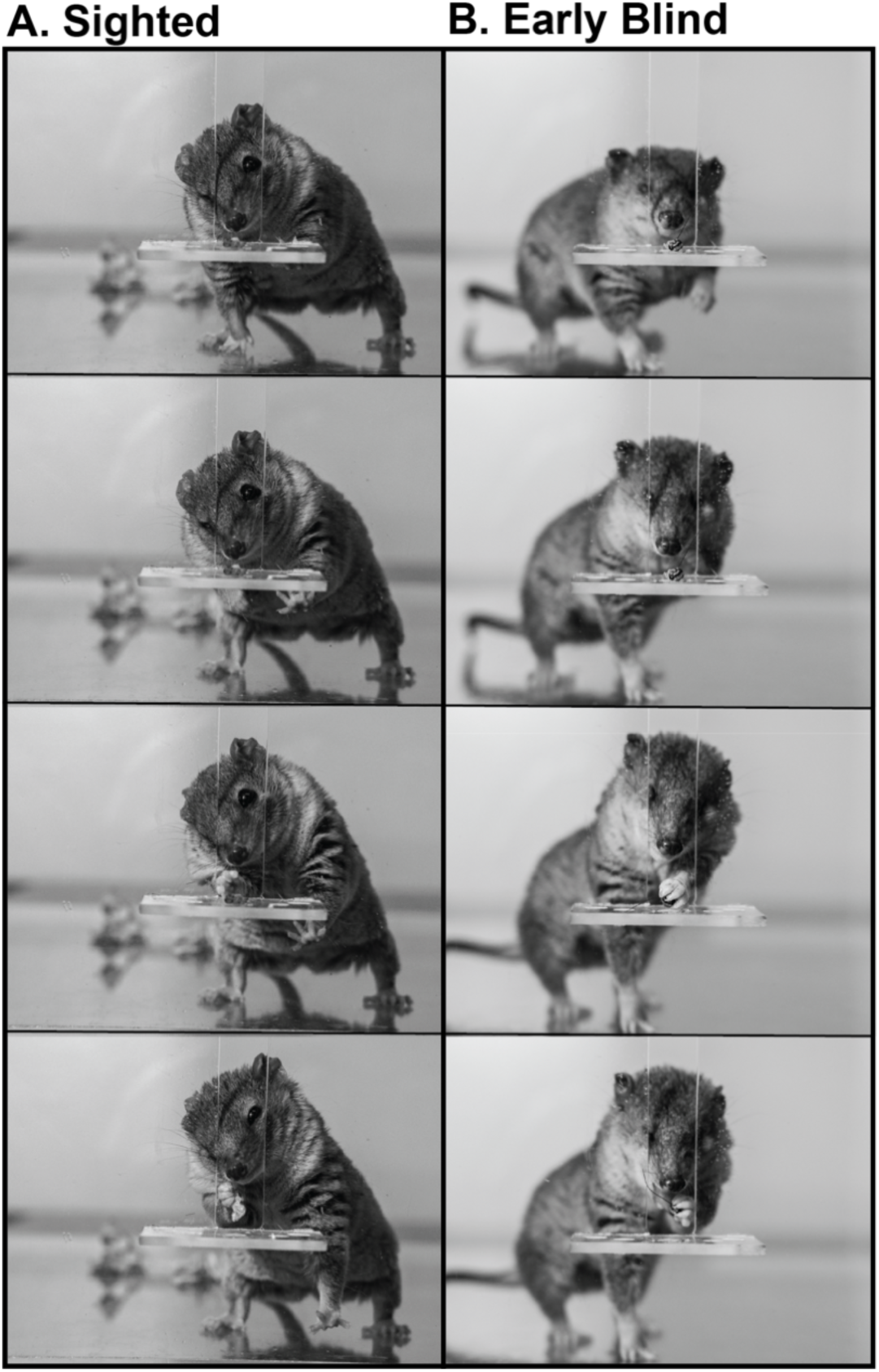
Images of reaching movements. **(A)** Individual video frames of a sighted control and **(B)** an early blind short-tailed opossums during the reaching and grasping task. Images of sighted and blind animals were matched according to phase of the task, starting with the initial reach (top images), then the successful grasp, and lastly bringing the food to the mouth (bottom images).

### Reaching and grasping in early blind and sighted opossums

Short-tailed opossums appeared to be innately adept at reaching and grasping, as both EB and SC animals rapidly achieved low-weighted error rates during shaping. Although SC animals appeared to exhibit a delayed reduction in weighted error across shaping days compared to EB animals, learning trajectories did not differ between visual groups across the 5-day shaping period (Figure S3A). No paw preference was observed in either group (Mann-Whitney U: 75.0, *p* = 0.169; Figure S3B). A linear mixed model showed a significant effect of biological sex on reach-to-grasp performance (Male vs Female: *p* = 0.0073; See Table 3 for EMM contrast; Figure S3C), with females committing more errors than males. These differences in weighted error by sex are a result of sexual dimorphism, where adult females, on average, are smaller than males (as indicated by lower body weight, smaller head size, and body length in females) (Bergallo and Cerqueira, 1994).

### Effects of vision on reaching performance

To determine whether visual cues facilitate reach-to-grasp performance under intact sensory conditions, we first analyzed baseline trials using a linear mixed-effects model with Visual Group (EB, SC), Lighting Condition (light, dark), Biological Sex, and a random intercept for animal identity (ID). Our model revealed a significant main effect of Lighting Condition (light, dark) (β = 15.23, SE = 5.76, t = 2.64, *p =* 0.013). Post-hoc estimated marginal means (EMM) analysis revealed that SC opossums committed significantly fewer errors in the light than in the dark (Table 3; Figure 4A). In contrast, EB opossums showed no significant effect of lighting and no significant differences in performance were detected between SC and EB animals under either light condition (Table 3; Figure 4A). These results suggest that visual cues substantially facilitate successful reach targeting for SC animals in this task, and that EB animals rely on other sensory modalities to compensate because vision is unavailable.

**Figure 4:**
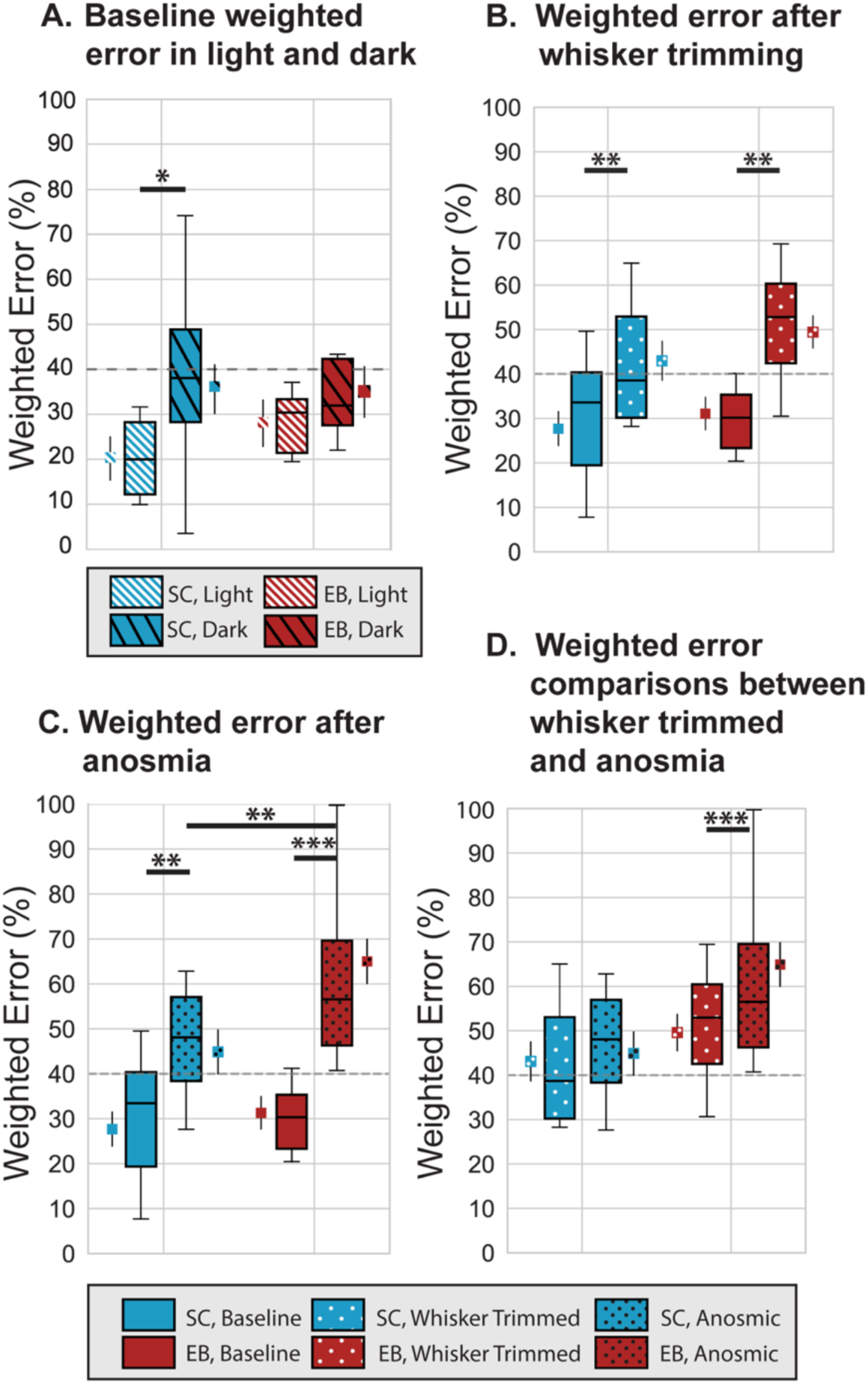
Reaching and grasping performance. Box and whisker plots show the distribution of animal-level mean percent weighted error for early blind animals (EB; red) and sighted controls (SC; blue). Estimated marginal means and standard errors are plotted as small boxes and whiskers adjacent to the main boxes. **(A)** SC opossums made significantly fewer errors in the light (white hatched) compared to the dark (black hatched) under baseline conditions, confirmed by estimated marginal means contrasts (EMM; small boxes beside whisker plots) (Light − Dark: p = 0.017). SC and EB opossums performed similarly under both light and dark conditions (Light: SC – EB, p = 0.28; Dark: SC – EB, p = 0.93). Additionally, EB animals showed no significant change in performance across lighting conditions (Light − Dark: p = 0.29). **(B)** Plots show performance before (solid) and after whisker trimming (white dotted) and indicate that the lack of tactile input significantly increased error in both groups (linear mixed model: *p* = 0.0016, not illustrated in figure), with estimated marginal means confirming significant increases in error for both EB and SC animals (Baseline – Trimmed: *p* = 0.0005) and sighted animals (Baseline – Trimmed: *p* = 0.0076). **(C)** Plots show performance before (solid) and after anosmia induction (black dotted). An increased error rate in both groups reflects the main effect of anosmia (linear mixed model: *p* = 0.0008, not illustrated in figure), with estimated marginal means confirming significant increased error in both sighted animals (Baseline – Anosmia: *p* = 0.0039) and EB animals (Baseline – Anosmia: p = 0.0001). A significant interaction between visual group (SC/EB) and pre-post anosmia was found (*p* = 0.031). EMM post-hoc analysis determined the direction of this effect: EB animals exhibited higher weighted error than sighted controls (SC/anosmic – EB/anosmic: *p* = 0.0055). **(D)** Plots comparing performance between whisker-trimmed (white dotted) and anosmic (black dotted) conditions. Early blind animals exhibited significantly greater targeting error under anosmia compared to whisker trimming (Anosmia – Trimmed: *p* = 0.0374). In contrast, sighted opossums showed no significant difference between the two conditions. For panels **B-D**, animal-level means were calculated across light and dark conditions. In all plots boxes represent interquartile range, with inside bar representing the median whiskers showing 1.5 times the interquartile range. ∗p < 0.05, ∗∗p < 0.01, ∗∗∗p < 0.001

### Reaching performance of early blind and sighted opossums is differentially affected by loss of input from other sensory systems

To examine the role of tactile input from the facial whiskers, and of olfaction in reach-to-grasp performance, we selectively restricted sensory input in these modalities in EB and SC opossums. Previous work from our laboratory has shown that EB opossums have alterations in receptive fields and response properties for neurons in the whisker representation of S1 and outperform SC animals in texture discrimination tasks (Ramamurthy et al, 2018, Ramamurthy et al, 2021; Figure 1C). Therefore, we trimmed the mystacial, genal, and submandibular vibrissae to determine their contributions to this reaching task in both SC and EB animals. As expected, we observed an overall increase in weighted error after whisker trimming in EB and SC animals (Figure S3D). Individual responses varied in magnitude; however, most animals showed increases in weighted error following whisker trimming, consistent with the group-level effects reported in Figure 4 (Figure S3F). Our linear mixed effects model revealed a significant positive effect of whisker trimming on targeting error rates of EB and SC opossums (β = 15.53, SE = 4.80, t = 3.238, *p* = 0.0016; Figure 4B) with no evidence for a differential effect between EB and SC animals (Table 3). Thus, whisker trimming had a similar impact on performance for both SC and EB opossums under both light and dark conditions, suggesting that information from the whiskers facilitates successful reach targeting in both groups.

To investigate the role of olfaction on reach-to-grasp, we induced anosmia and tested reach-to-grasp performance under light and dark conditions. Similar to whisker trimming, we observed an overall increase in average weighted error when comparing performance before and after anosmia induction (Figure S3E). Individual responses varied in magnitude; however, most animals showed increased weighted error after anosmia induction, consistent with the group-level effects reported in Figure 4 (Figure S3F). Our linear mixed-effects model then revealed a significant increase in weighted error following anosmia induction across both visual groups (β = 18.03, SE = 5.20, t = 3.47, *p =* 0.0008; Figure 4C). Importantly, there was a significant interaction effect between Visual Group (EB /SC) and Sensory Condition (baseline/anosmic) (β = 16.40, SE = 7.50, t = 2.19, *p* = 0.031), indicating that the effect of anosmia on target error rate differed between EB and SC animals. Post-hoc estimated marginal means revealed that under anosmia, EB animals exhibited significantly larger weighted error rates than SC animals (Table 3; Figure 4C). Thus, anosmia disproportionately reduced EB performance, suggesting that olfactory information substantially facilitates successful reach targeting for EB animals.

When comparing performance between anosmia and whisker-trimmed sensory manipulations, EB animals exhibited significantly greater targeting errors under anosmia compared to whisker trimming across lighting conditions (Table 3; Figure 4D), suggesting that EB opossums rely more on olfactory input than tactile input from the vibrissae. In contrast, although the performance of SC opossums was affected by loss of tactile input and olfactory input, there was no significant difference in their performance between the two manipulations (Table 3; Figure 4D), underscoring that their performance remains more stable across different sensory disruptions. These findings suggest that EB animals may rely more heavily on olfactory input to guide forelimb movements, whereas SC animals appear less affected by its loss, potentially relying more on visual cues.

## Discussion

We examined how early vision loss and subsequent tactile or olfactory deprivation affected reach-to-grasp performance in adult short-tailed opossums. Early blind (EB) short tailed opossums performed accurate reach-to-grasp movements, suggesting that the remaining senses can compensate for vision loss. Selective removal of tactile and olfactory input revealed that both EB and SC animals relied on these sensory modalities to retrieve prey, with olfaction and whisker touch contributing strongly to reach accuracy in EB animals.

### Sensory-guided reaching and grasping in non-primate models

Primates are considered the reaching and grasping experts among mammals, especially those with opposable thumbs such as macaques, capuchin monkeys and humans (Christel and Fragaszy, 2000; Fox et al., 2019; Mitchell et al., 2024). Despite this emphasis on primates, most reach-and-grasp studies use mice and rats (Whishaw et al., 2017; Naghizadeh et al., 2020; Parmiani et al., 2021; Quarta et al., 2022), and our results demonstrate that opossums are also innately skilled at reaching and grasping (Figure S3A).

Although little is known about the predatory behavior of short-tailed opossums, laboratory studies suggest that they rely heavily on forepaw capture, while rats capture prey with the mouth (Ivanco et al., 1996). Comparisons of naïve reaching strategies show that opossums are more precise, achieving higher success rates on first attempts, while rats appear less coordinated and require more training to achieve comparable success (Ivanco et al., 1996). Given that locomotor habit exerts strong selective pressure on limb morphology, these differences between opossums and rats may reflect the semi-arboreal niche that short-tailed opossums occupy in the wild compared to the terrestrial niche that brown rats would naturally occupy. Short-tailed opossums possess arboreal adaptations that likely enhance forepaw dexterity (Ivanco et al., 1996), suggesting that selection for climbing may also favor single paw reaching. Consistent with this idea, studies on arboreal (*T. minor)* and terrestrial (*T. tana*) tree shrews reveal large differences in manual dexterity and reaching that correlate with the degree of habitual arboreal locomotion. Specifically, *T. minor* consistently relied on single-paw grasping strategies, while *T. tana* was never observed grasping food rewards with the paws (Sargis, 2001). These comparisons indicate that adaptations to an arboreal lifestyle coincide with selection for single paw reaching and grasping capacities. These interspecies differences highlight the importance of species-specific motor and sensory adaptations when evaluating reach-to-grasp capabilities, making the short-tailed opossum a suitable single-paw model for understanding reaching behaviors and their underlying neural circuits.

### The contribution of different senses to reaching and grasping

All mammals guide their sensory effectors to control information sampling during exploration. These movements are not fixed but are dynamically adjusted according to task demands, allowing animals to adopt sampling strategies that maximize reward outcomes. For example, whisking movement strategies are modulated during the exploration of a novel object (Gordon et al., 2014). This framework provides the foundation for examining how short-tailed opossums utilize whisker touch, olfaction, and vision during reaching and grasping, and how the relative roles of these senses shift after early vision loss.

Our results demonstrate that vision facilitates reaching in short-tailed opossums, as SC animals committed more targeting errors in the dark than in the light (Figure 4A). Nevertheless, SC opossums still performed the task in darkness, suggesting that although vision is beneficial, it is not necessary for success. Instead, nonvisual sensory cues contributed substantially to performance in SC animals and likely served as the primary source of sensory guidance in EB animals.

Studies from our laboratory found that EB opossums outperformed SC opossums in ladder-rung walking tasks, and compensated via tactile sensing with the mystacial whiskers, since trimming these whiskers decreased task performance (Englund et al., 2020). A subsequent study demonstrated that EB opossums had lower texture discrimination thresholds compared to SC opossums, an effect that disappeared after whisker trimming (Ramamurthy et al., 2021). In the current study, whisker trimming increased targeting error in both EB and SC animals, indicating that tactile input contributes to reaching performance and is a key driver of behavioral compensation following early loss of vision (Figure 4B).

In EB opossums, greater reliance on whisker touch coincides with smaller receptive fields and sharper tuning of neurons in the whisker representation of S1 (Ramamurthy and Krubitzer, 2018). Evidence also suggests that while the basic pattern of connections of V1 is still present in EB animals, the density of these connections changes and novel cortical and subcortical inputs to V1 from somatosensory neocortex and thalamic nuclei emerge (Karlen et al., 2006). Similarly, in anophthalmic mice, differences in functional, and anatomical organization of the brain are observed (Chabot et al., 2007, 2008; Massé et al., 2014). However, reduced connectivity between S1 and “V1 in anophthalmic mice together with changes in the density of pre-existing inputs from other cortical areas (Charbonneau et al., 2012), suggests selective strengthening and weakening of pre-existing inputs to “V1”, consistent with mechanisms described following sensory deprivation (Ewall et al., 2021).

Studies in EB humans also demonstrate that they have enhanced tactile acuity and vibrotactile perception (Wan et al., 2010; Radziun et al., 2023), potentially facilitated by more precise spatial tuning of tactile attention in EB versus sighted individuals (Forster et al., 2007). Whether these enhancements arise from early blindness itself or from altered tactile experience is unclear. Enhanced passive tactile acuity can occur independently of childhood visual experience, light perception, or Braille reading (Goldreich and Kanics, 2003), whereas differences between Braille readers and non-readers suggest that tactile experience can further enhance acuity (Wong et al., 2011b). These findings indicate that both early sensory deprivation and experience-dependent plasticity contribute to enhanced tactile abilities in congenital blindness, although their relative contributions are not well understood.

EB humans exhibit functional differences in brain organization as well as behavioral differences. V1, extrastriate cortex, and posterior parietal cortex are activated during Braille reading (Sadato et al., 1996; Buchel, 1998), implicating these regions in tactile discrimination and language processing. Particularly relevant to the current study, early visual areas (V1, V2 and V3), encode reach direction in EB individuals, together with other visuospatial and visuomotor cortical regions (Bola et al., 2023). However, cortical areas beyond early visual cortex may not undergo extensive functional reorganization following early blindness. The human middle temporal cortex (hMT+), which contributes to motion and optic flow processing in sighted individuals, is recruited during tactile flow in EB individuals, suggesting that computations underlying self-motion perception can operate independently of a specific sensory modality (Ricciardi et al., 2007; Leo et al., 2012). Similarly, sensory-guided hand movements activate the superior parietal cortex (SPC) of EB and sighted individuals, suggesting that computations supporting complex manual behaviors operate independently of the sensory modality guiding hand movements (Fiehler et al., 2009). Based on accumulating evidence, Ricciardi and Pietrini (2024) propose that regions such as SPC and hMT+ perform modality-invariant computations, preserving aspects of their functional role despite differences in sensory input. Whether recruitment of V1 during higher-order cognitive tasks similarly reflects modality-independent computations or more extensive functional reorganization is also an open question (Ricciardi et al., 2020; Makin and Krakauer, 2023; Saccone et al., 2024).

Although the olfactory capabilities of short-tailed opossums remain uncharacterized, this study demonstrates that both EB and SC opossums rely on olfaction to target prey, and that this sense is critical in EB animals (Figure 4C, D). This is unsurprising because smell is the only way EB opossums can locate immobile prey beyond whisker range. Similarly, anophthalmic mice outperform sighted mice in olfactory tasks such as retrieving buried food, and these differences coincide with increased volume of olfactory structures like the piriform cortex, the amygdala, and the glomerular and granular layers of the olfactory bulb (Touj et al., 2020, 2021). Congenitally blind humans demonstrate lower odor detection thresholds (Beaulieu-Lefebvre et al., 2011), increased sensitivity to odorants (Gagnon et al., 2014), and better performance on odorant localization tasks (Manescu et al., 2020) than sighted individuals. Functional imaging studies show that these behavioral differences coincide with stronger activation of olfactory structures (e.g., orbitofrontal cortex, amygdala, and hippocampus) as well as extensive activation of occipital cortex during task performance (Kupers et al., 2011).

Taken together, data from congenitally blind humans and other mammals demonstrate differences in tactile and olfactory behaviors. Reaching and grasping requires representations of peripersonal space and the ability to distinguish self from the surrounding environment (Ricciardi et al., 2007, 2017), processes that, in sighted individuals, rely on visual, tactile, and proprioceptive information. Although direct comparisons between humans and opossums are limited by species differences in visual dependence, and evidence for supramodal organization in non-primate models of early blindness remains scarce, the ability of both SC and EB opossums to successfully reach and grasp is consistent with the possibility that computations supporting goal-direct actions can operate using different sources of sensory information. This study suggests that the computations supporting goal-directed actions in peripersonal space are preserved despite the absence of vision, through stable supramodal computations and experience-dependent plasticity within spared sensory systems that enables remaining sensory modalities to guide reaching and grasping.

## Conflict of interest statement

The authors declare no conflict of interest.

## Acknowledgments

We thank Chris S. Bresee and Fernando Gomez for feedback on statistical analysis and for help with editing the final manuscript. We thank Phoenix Twomey for help with animal care and illustrations. This research was funded by NIH NEI Grants 5R01EY034303 to L.A.K., F31EY034400 to C.R.P., and T32 EY015387.

## Author contributions

L.A.K. and C.R.P. designed research; C.R.P., F.Z.N., M.M. performed behavioral testing; C.R.P. data curation; C.R.P. analyzed data; C.R.P. data visualization; C.R.P. manuscript preparation; L.A.K. and C.R.P. manuscript editing; L.A.K. supervision; L.A.K. resources.

---

**Supplementary Figure 1:**
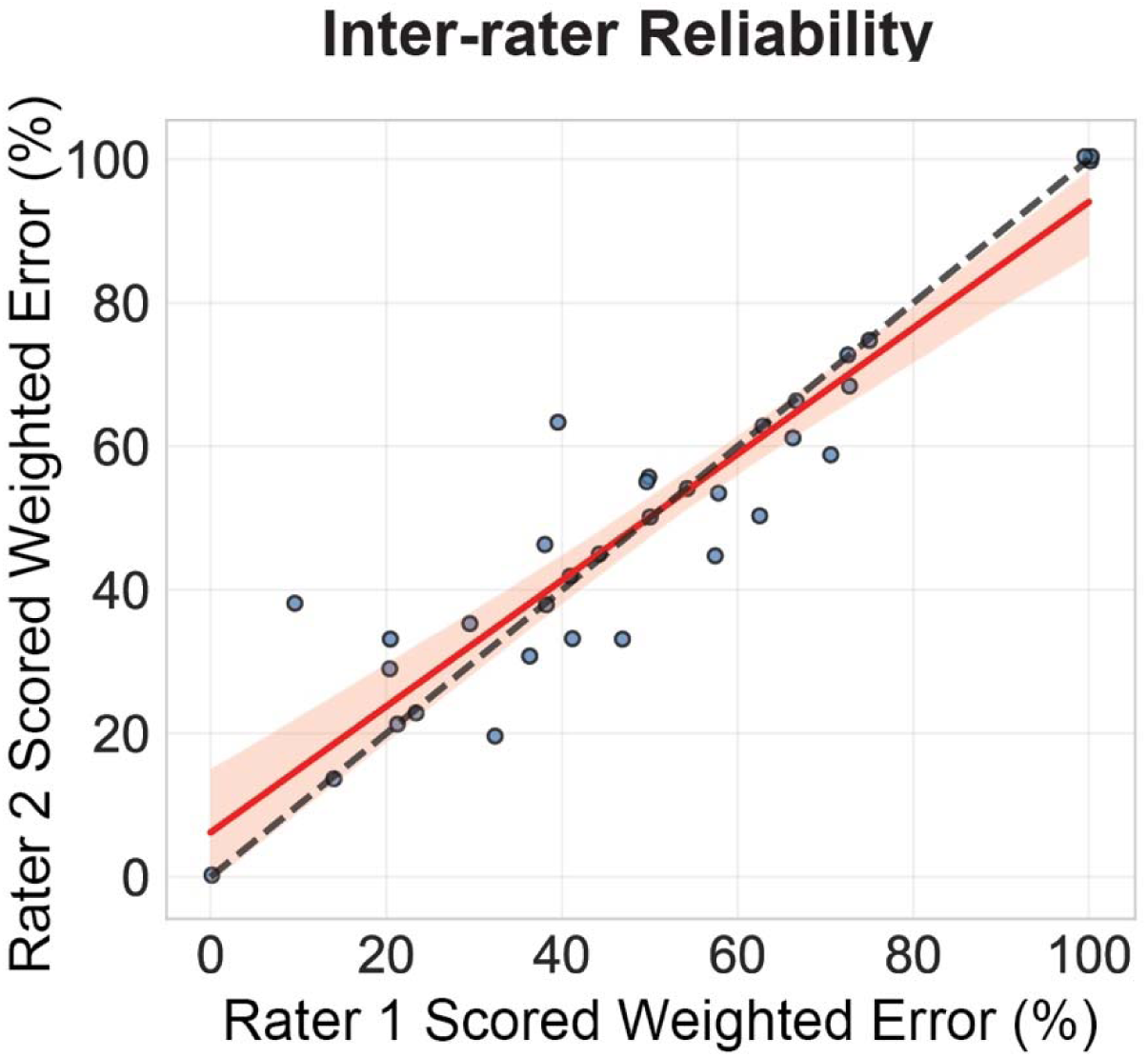
Scatterplot with regression line (red) and line of perfect agreement (dashed) showing the degree of agreement between raters on weighted error scoring. The fitted regression line indicated strong correspondence between raters (R²adj = 0.87, F(1,32) = 218.0, p < 0.001). Interrater reliability, assessed using intraclass correlation (ICC3, single fixed raters), revealed excellent agreement (ICC3= 0.96, p < .001).

**Supplementary Figure 2:**
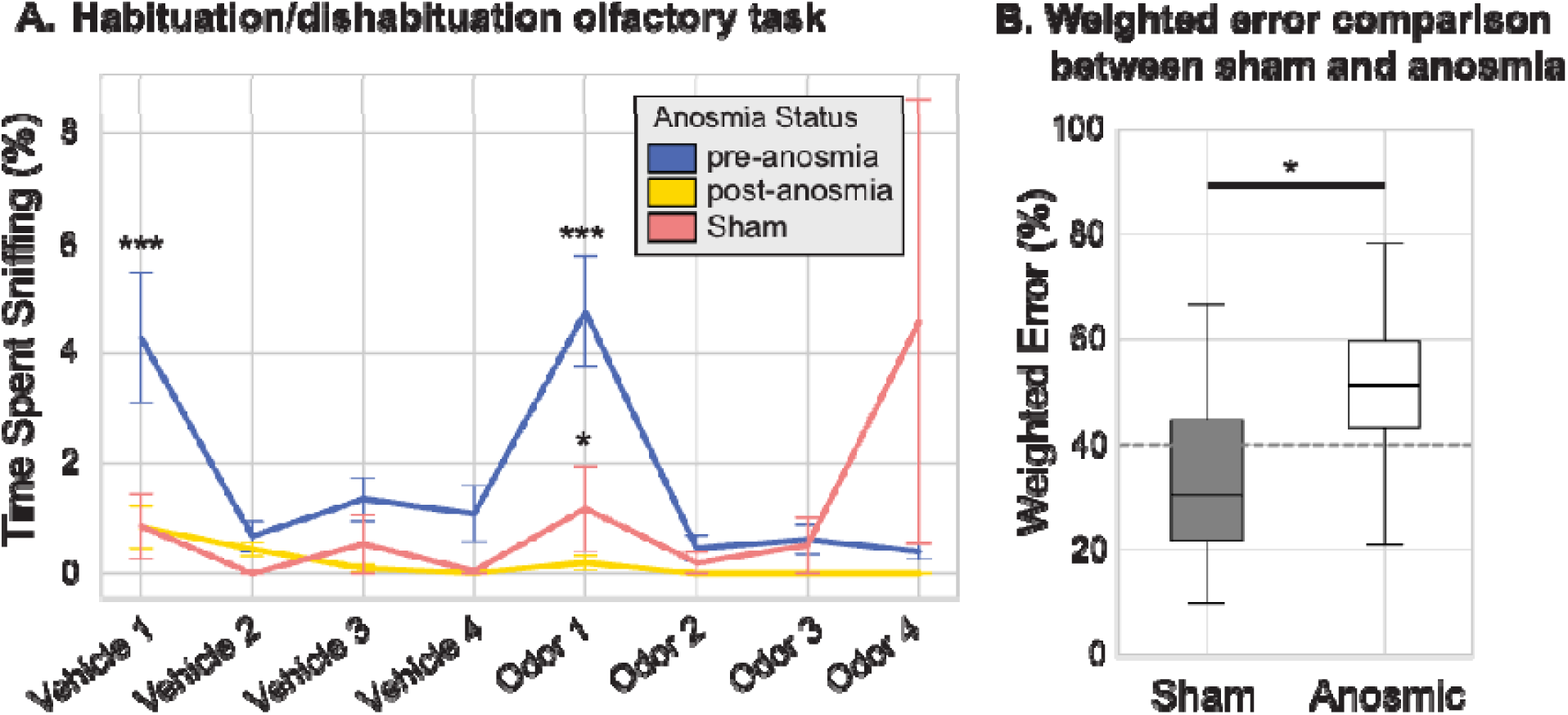
**(A)** Line graph shows average percent sniffing across odor presentations before (blue), after anosmia induction (yellow) and after sham procedures (pink). These tests were performed in the light. Error bars represent bootstrapped 95% confidence intervals. Anosmia induction significantly decreased the average time spent sniffing (Odor 1: *p* = 0.012) in both groups of animals. We observed a significant difference in sniffing time for the first presentation of the vehicle (Vehicle 1: *p* = 0.01), as well as the first presentation of the odor in non-anosmic animals. This difference disappeared following anosmia. Sham-treated opossums retained odor-directed investigation behavior and spent significantly more time attending to odor stimuli than post-anosmic animals (*p* = 0.024). **(B)** Box and whisker plots show sham (grey) and anosmic (white) reaching and grasping errors averaged across blind and sighted animals and lighting conditions. Anosmia significantly increased weighted error rates compared to sham inductions (*p* = 0.024). Notations are the same as in the previous figures; (∗*p* < 0.05, ∗∗*p* < 0.01, ∗∗∗*p* < 0.001).

**Supplementary Figure 3:**
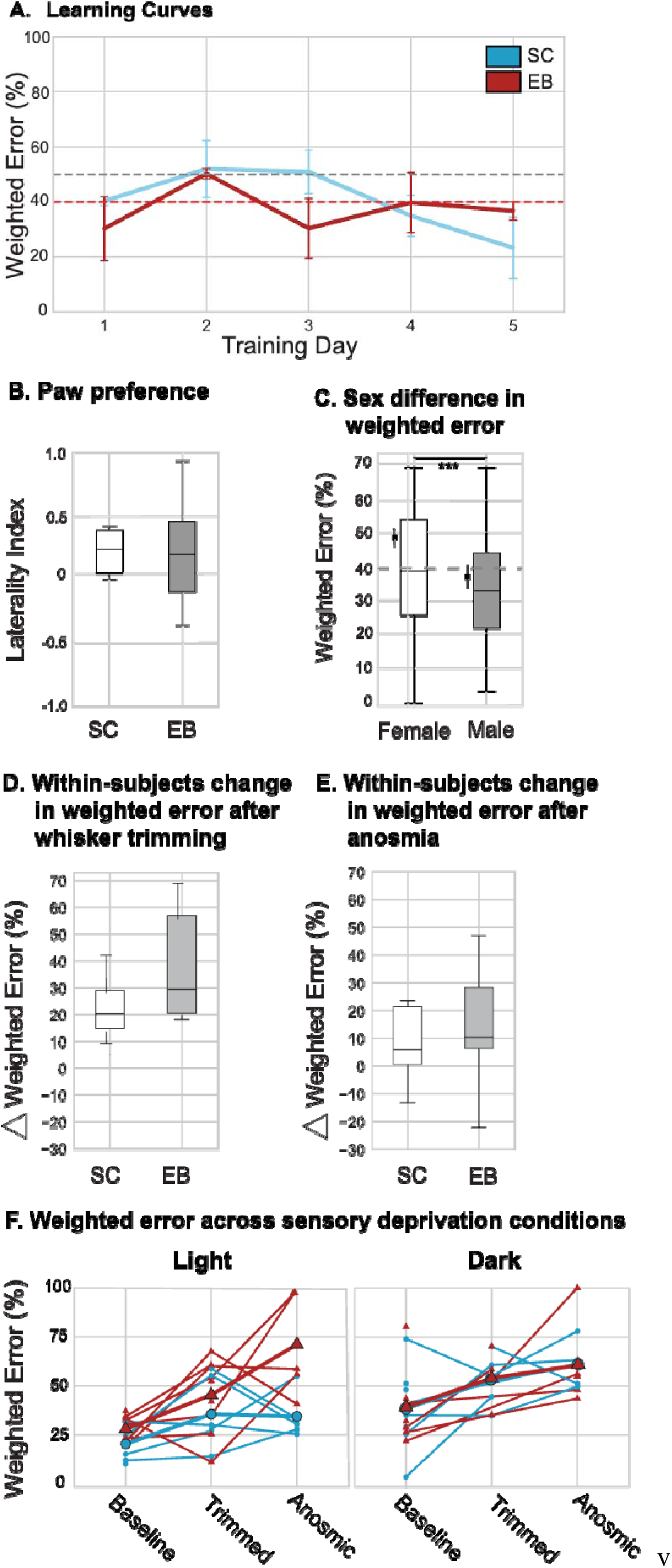
**(A)** Weighted error rates (%) across five consecutive training days for sighted (SC; blue) and early blind (EB; red) short-tailed opossums. Data points represent group means and whiskers denote the standard error of the mean (SEM). Dashed horizontal lines indicate training criteria (grey = 50% chance performance; red = 40% criterion threshold). **(B)** Box and whisker plots show that EB and sighted short-tailed opossums do not exhibit significant differences in paw preference (Mann-Whitney U = 75.0, *p* = 0.169). **(C)** Box and whisker plots illustrate percent weighted error for female (white) and male (gray) opossums across all visual groups, sensory conditions, and light conditions. A linear mixed model revealed a significant main effect of sex, with females committing significantly more targeting errors than males (β = – 11.58 ± 3.68 SE, t = – 3.14, *p* = 0.0073). Estimated marginal means indicated that females had higher error rates compared to males (Female – Male: *p* = 0.012). Estimated marginal means and standard errors are plotted as small boxes and whiskers adjacent to the main boxes. The dashed line marks the 40% weighted error criterion. **(D)** Within-subjects average change in weighted error rate before and after whisker trimming. Positive values indicate overall increases in weighted error rates across all animals, although the extent of the effect varied. **(E)** Within-subjects average change in weighted error rate before and after anosmia induction. Positive values indicate overall increases in weighted error rates in SC and EB animals, although the effect was greater in EB animals. Boxes represent the middle 50% of the data (interquartile range); the black horizontal line indicates the median; whiskers extend to 1.5 times the interquartile range. **(F)** Weighted error trajectories across sensory deprivation conditions. Individual animal performance is shown for the baseline, whisker-trimmed, and anosmia conditions under light (left) and dark (right) testing conditions. Each colored line represents a single animal with sighted control (SC; blue) and early blind (EB; red) opossums shown separately, illustrating changes in mean weighted error across sensory manipulations. Animals generally exhibited increased weighted error following sensory deprivation, although the magnitude of the effect varied across individuals. Values represent the mean weighted error (%) for each animal within each sensory condition.

## Notes

### Competing Interest Statement

The authors have declared no competing interest.

### Summary of Updates

The revision includes a revised Figure 4 that presents animal-level performance data, including a revised Figure 4A and a new corresponding analysis of baseline performance across light and dark conditions; a new table reporting animal-level session contributions across testing conditions; a new panel in Supplementary Figure 3 showing individual animal performance across all sensory and lighting conditions; and a substantially revised Discussion that incorporates a more balanced account of evidence from the human literature on early blindness.

## References

Alaverdashvili M, Whishaw IQ (2013) A behavioral method for identifying recovery and compensation: Hand use in a preclinical stroke model using the single pellet reaching task. Neuroscience & Biobehavioral Reviews 37:950–967.

Asumbisa K, Peyrache A, Trenholm S (2022) Flexible cue anchoring strategies enable stable head direction coding in both sighted and blind animals. Nat Commun 13:5483.

Beaulieu-Lefebvre M, Schneider FC, Kupers R, Ptito M (2011) Odor perception and odor awareness in congenital blindness. Brain Research Bulletin 84:206–209.

Bergallo HG, Cerqueira R (1994) Reproduction and growth of the opossum Monodelphis domestica (Mammalia: Didelphidae) in northeastern Brazil. Journal of Zoology 232:551–563.

Bola Ł, Vetter P, Wenger M, Amedi A (2023) Decoding Reach Direction in Early “Visual” Cortex of Congenitally Blind Individuals. J Neurosci 43:7868–7878.

Buchel C (1998) Different activation patterns in the visual cortex of late and congenitally blind subjects. Brain 121:409–419.

Chabot N, Charbonneau V, Laramée M-E, Tremblay R, Boire D, Bronchti G (2008) Subcortical auditory input to the primary visual cortex in anophthalmic mice. Neuroscience Letters 433:129–134.

Chabot N, Robert S, Tremblay R, Miceli D, Boire D, Bronchti G (2007) Audition differently activates the visual system in neonatally enucleated mice compared with anophthalmic mutants. European Journal of Neuroscience 26:2334–2348.

Charbonneau V, Laramée M-E, Boucher V, Bronchti G, Boire D (2012) Cortical and subcortical projections to primary visual cortex in anophthalmic, enucleated and sighted mice. European Journal of Neuroscience 36:2949–2963.

Christel MI, Fragaszy D (2000) Manual Function in Cebus apella. Digital Mobility, Preshaping, and Endurance in Repetitive Grasping. International Journal of Primatology 21:697–719.

Englund M, Faridjoo S, Iyer CS, Krubitzer L (2020) Available Sensory Input Determines Motor Performance and Strategy in Early Blind and Sighted Short-Tailed Opossums. iScience 23:101527.

Ewall G, Parkins S, Lin A, Jaoui Y, Lee H-K (2021) Cortical and Subcortical Circuits for Cross-Modal Plasticity Induced by Loss of Vision. Front Neural Circuits 15:665009.

Fiehler K, Burke M, Bien S, Röder B, Rösler F (2009) The Human Dorsal Action Control System Develops in the Absence of Vision. Cereb Cortex 19:1–12.

Forster B, Eardley AF, Eimer M (2007) Altered tactile spatial attention in the early blind. Brain Research 1131:149–154.

Fox DM, Mundinano I-C, Bourne JA (2019) Prehensile kinematics of the marmoset monkey: Implications for the evolution of visually-guided behaviors. Journal of Comparative Neurology 527:1495–1507.

Friard O, Gamba M (2016) BORIS: a free, versatile open-source event-logging software for video/audio coding and live observations. Methods in Ecology and Evolution 7:1325–1330.

Fujii T, Tanabe HC, Kochiyama T, Sadato N (2009) An investigation of cross-modal plasticity of effective connectivity in the blind by dynamic causal modeling of functional MRI data. Neuroscience Research 65:175–186.

Gagnon L, Kupers R, Ptito M (2014) Making Sense of the Chemical Senses. Available at: https://brill.com/view/journals/msr/27/5-6/article-p399_8.xml [Accessed April 21, 2026].

Goldreich D, Kanics IM (2003) Tactile Acuity is Enhanced in Blindness. J Neurosci 23:3439–3445.

Gordon G, Fonio E, Ahissar E (2014) Emergent Exploration via Novelty Management. J Neurosci 34:12646–12661.

Hagan MA, Rosa MGP, Lui LL (2017) Neural plasticity following lesions of the primate occipital lobe: The marmoset as an animal model for studies of blindsight. Developmental Neurobiology 77:314–327.

Hunt DM, Chan J, Carvalho LS, Hokoc JN, Ferguson MC, Arrese CA, Beazley LD (2009) Cone visual pigments in two species of South American marsupials. Gene 433:50–55.

Ivanco TL, Pellis SM, Whishaw IQ (1996) Skilled forelimb movements in prey catching and in reaching by rats (*Rattus norvegicus*) and opossums (*Monodelphis domestica*): relations to anatomical differences in motor systems. Behavioural Brain Research 79:163–181.

Izraeli R, Koay G, Lamish M, Heicklen-Klein AJ, Heffner HE, Heffner RS, Wollberg Z (2002) Cross-modal neuroplasticity in neonatally enucleated hamsters: structure, electrophysiology and behaviour. European Journal of Neuroscience 15:693–712.

Kahn DM, Krubitzer L (2002) Massive cross-modal cortical plasticity and the emergence of a new cortical area in developmentally blind mammals. Proceedings of the National Academy of Sciences 99:11429–11434.

Karlen SJ, Kahn DM, Krubitzer L (2006) Early blindness results in abnormal corticocortical and thalamocortical connections. Neuroscience 142:843–858.

Klein A, Sacrey L-AR, Whishaw IQ, Dunnett SB (2012) The use of rodent skilled reaching as a translational model for investigating brain damage and disease. Neuroscience & Biobehavioral Reviews 36:1030–1042.

Kupers R, Beaulieu-Lefebvre M, Schneider FC, Kassuba T, Paulson OB, Siebner HR, Ptito M (2011) Neural correlates of olfactory processing in congenital blindness.Neuropsychologia 49:2037–2044.

Leo A, Bernardi G, Handjaras G, Bonino D, Ricciardi E, Pietrini P (2012) Increased BOLD Variability in the Parietal Cortex and Enhanced Parieto-Occipital Connectivity during Tactile Perception in Congenitally Blind Individuals. Neural Plasticity 2012:720278.

Makin TR, Krakauer JW (2023) Against cortical reorganisation Pruszynski JA, de Lange FP, eds. eLife 12:e84716.

Manescu S, Chouinard-Leclaire C, Collignon O, Lepore F, Frasnelli J (2020) Enhanced Odorant Localization Abilities in Congenitally Blind but not in Late-Blind Individuals. Chem Senses 46:bjaa073.

Maruoka S, Sugano E, Togawa R, Katayama N, Tabata K, Yoshizawa N, Morita R, Takita Y, Ozaki T, Fukuda T, Bai L, Tomita H (2024) Enhanced auditory responses in visual cortex of blind rats using intrinsic optical signal imaging. Sci Rep 14:24740.

Massé IO, Guillemette S, Laramée M-E, Bronchti G, Boire D (2014) Strain differences of the effect of enucleation and anophthalmia on the size and growth of sensory cortices in mice. Brain Research 1588:113–126.

Metz GA, Whishaw IQ (2002) Cortical and subcortical lesions impair skilled walking in the ladder rung walking test: a new task to evaluate fore- and hindlimb stepping, placing, and co-ordination. Journal of Neuroscience Methods 115:169–179.

Mitchell JF, Wang KH, Batista AP, Miller CT (2024) An ethologically motivated neurobiology of primate visually-guided reach-to-grasp behavior. Current Opinion in Neurobiology 86:102872.

Naghizadeh M, Mohajerani MH, Whishaw IQ (2020) Mouse Arm and hand movements in grooming are reaching movements: Evolution of reaching, handedness, and the thumbnail. Behavioural Brain Research 393:112732.

Parmiani P, Lucchetti C, Franchi G (2021) Changes in reach-to-grasp behaviour over the course of training in rats. European Journal of Neuroscience 54:7805–7819.

Quarta E, Scaglione A, Lucchesi J, Sacconi L, Allegra Mascaro AL, Pavone FS (2022) Distributed and Localized Dynamics Emerge in the Mouse Neocortex during Reach-to-Grasp Behavior. J Neurosci 42:777–788.

Radziun D, Crucianelli L, Korczyk M, Szwed M, Ehrsson HH (2023) The perception of affective and discriminative touch in blind individuals. Behavioural Brain Research 444:114361.

Ramamurthy DL, Dodson HK, Krubitzer LA (2021) Developmental plasticity of texture discrimination following early vision loss in the marsupial *Monodelphis domestica*. Journal of Experimental Biology 224:jeb236646.

Ramamurthy DL, Krubitzer LA (2018) Neural Coding of Whisker-Mediated Touch in Primary Somatosensory Cortex Is Altered Following Early Blindness. J Neurosci 38:6172–6189.

Ricciardi E, Menicagli D, Leo A, Costantini M, Pietrini P, Sinigaglia C (2017) Peripersonal space representation develops independently from visual experience. Sci Rep 7:17673.

Ricciardi E, Papale P, Cecchetti L, Pietrini P (2020) Does (lack of) sight matter for V1? New light from the study of the blind brain. Neuroscience & Biobehavioral Reviews 118:1–2.

Ricciardi E, Pietrini P (2024) The supramodality “spillover” from neuroscience to cognitive sciences: a commentary on Calzavarini (2024). Language, Cognition and Neuroscience 39:867–871.

Ricciardi E, Vanello N, Sani L, Gentili C, Scilingo EP, Landini L, Guazzelli M, Bicchi A, Haxby JV, Pietrini P (2007) The Effect of Visual Experience on the Development of Functional Architecture in hMT+. Cerebral Cortex 17:2933–2939.

Saccone EJ, Tian M, Bedny M (2024) Developing cortex is functionally pluripotent: Evidence from blindness. Developmental Cognitive Neuroscience 66:101360.

Sadato N, Pascual-Leone A, Grafman J, Ibañez V, Deiber M-P, Dold G, Hallett M (1996) Activation of the primary visual cortex by Braille reading in blind subjects. Nature 380:526–528.

Sargis EJ (2001) The grasping behaviour, locomotion and substrate use of the tree shrews Tupaia minor and T. tana (Mammalia, Scandentia). Journal of Zoology 253:485–490.

Seabold S, Perktold J (2010) Statsmodels: Econometric and Statistical Modeling with Python. Proceedings of the 9th Python in Science Conference 2010.

Setti F, Handjaras G, Bottari D, Leo A, Diano M, Bruno V, Tinti C, Cecchetti L, Garbarini F, Pietrini P, Ricciardi E (2023) A modality-independent proto-organization of human multisensory areas. Nat Hum Behav 7:397–410.

Sharma J, Angelucci A, Sur M (2000) Induction of visual orientation modules in auditory cortex. Nature 404:841–847.

Stone KD, Gonzalez CLR (2015) The contributions of vision and haptics to reaching and grasping. Front Psychol 6:1403.

Touj S, Cloutier S, Jemâa A, Piché M, Bronchti G, Al Aïn S (2020) Better Olfactory Performance and Larger Olfactory Bulbs in a Mouse Model of Congenital Blindness. Chem Senses 45:523–531.

Touj S, Gallino D, Chakravarty MM, Bronchti G, Piché M (2021) Structural brain plasticity induced by early blindness. European Journal of Neuroscience 53:778–795.

Voller J, Potužáková B, Šimeček V, Vožeh F (2014) The role of whiskers in compensation of visual deficit in a mouse model of retinal degeneration. Neuroscience Letters 558:149–153.

Wan CY, Wood AG, Reutens DC, Wilson SJ (2010) Congenital blindness leads to enhanced vibrotactile perception. Neuropsychologia 48:631–635.

Wang R, Wu L, Tang Z, Sun X, Feng X, Tang W, Qian W, Wang J, Jin L, Zhong Y, Xiao Z (2017) Visual cortex and auditory cortex activation in early binocularly blind macaques: A BOLD-fMRI study using auditory stimuli. Biochemical and Biophysical Research Communications 485:796–801.

Whishaw IQ (1996) An endpoint, descriptive, and kinematic comparison of skilled reaching in mice (Mus musculus) with rats (Rattus norvegicus). Behavioural Brain Research 78:101–111.

Whishaw IQ, Faraji J, Kuntz JR, Mirza Agha B, Metz GAS, Mohajerani MH (2017) The syntactic organization of pasta-eating and the structure of reach movements in the head-fixed mouse. Sci Rep 7:10987.

Whishaw IQ, Travis SG, Koppe SW, Sacrey L-A, Gholamrezaei G, Gorny B (2010) Hand shaping in the rat: Conserved release and collection vs. flexible manipulation in overground walking, ladder rung walking, cylinder exploration, and skilled reaching. Behavioural Brain Research 206:21–31.

Wong M, Gnanakumaran V, Goldreich D (2011a) Tactile Spatial Acuity Enhancement in Blindness: Evidence for Experience-Dependent Mechanisms. J Neurosci 31:7028–7037.

Wong M, Gnanakumaran V, Goldreich D (2011b) Tactile Spatial Acuity Enhancement in Blindness: Evidence for Experience-Dependent Mechanisms. J Neurosci 31:7028–7037.

